# Amino and PEG-Amino Graphene Oxide Grids Enrich and Protect Samples for High-resolution Single Particle Cryo-electron Microscopy

**DOI:** 10.1101/813972

**Authors:** Feng Wang, Zanlin Yu, Miguel Betegon, Melody Campbell, Tural Aksel, Jianhua Zhao, Sam Li, Shawn M. Douglas, Yifan Cheng, David A. Agard

## Abstract

Cryo-EM samples prepared using the traditional methods often suffer from too few particles, poor particle distribution, or strongly biased orientation, or damage from the air-water interface. Here we report that functionalization of graphene oxide (GO) coated grids with amino groups concentrates samples on the grid with improved distribution and orientation. By introducing a PEG spacer, particles are kept away from both the GO surface and the air-water interface, protecting them from potential denaturation.

## 1. Introduction

Single particle cryo-electron microscopy (cryo-EM) has become a major method for determining protein structures at a near-atomic or atomic resolution(Cheng, 2015; Nogales, 2016). Despite rapid advances in image collection and processing, sample preparation has remained largely unchanged, and can be rate limiting. In the standard cryo-EM specimen preparation process, the protein sample is applied onto a holey cryo-EM grid and the grid is plunged into liquid ethane after blotting to obtain a thin layer of vitrified amorphous ice, in principle, preserving the biological sample in a hydrated state (Mcdowall et al., 1983; Taylor and Glaeser, 1974). In practice, however, there are many issues preventing a sample from being in an ideal state for data collection. First, some particles strongly prefer to stick to the carbon film instead of being visible within the holes(Snijder et al., 2017). Second, some proteins cannot be concentrated to a sufficient level for cryo-EM sampling. Third, proteins or complexes may adopt “preferred orientations”, dissociate or denature as a result of interaction with the air-water interface(D’Imprima et al., 2019; Tan et al., 2017). This is now understood to be a much more pervasive problem than originally thought (Noble et al., 2018). Numerous approaches have been aimed at combatting these problems, including PEGylation on gold coated holey carbon grids(Meyerson et al., 2014), addition of detergents (Chen et al., 2019), use of grids deposited with a thin continuous carbon film(Nguyen et al., 2015) or surface modified graphene as substrate(D’Imprima et al., 2019; Naydenova et al., 2019), collecting data at defined tilts(Tan et al., 2017) and so on. However, no single strategy has yet to be widely adopted due to technical challenges, cost of materials, requirements for highly specialized equipment to make single crystal graphene films, or the deterioration of signal-to-noise and resolution experienced with thin carbon films.

To circumvent these problems, we first developed a simple and convenient approach to efficiently deposit graphene oxide (GO) films coated EM-grids(Palovcak et al., 2018), and have since explored various strategies for chemical functionalization. Here we discuss the broad utility of amino-functionalized GO grids (**Figure 1A**). The resulting grids provided a robust and broadly useful affinity to proteins without selectivity. We tested four different samples on our grids and found that 1)proteins were significantly enriched, much like on negative stained carbon grids, 2) particles were nicely distributed over grid holes 3) particle orientations were altered, without being overly biased. Using the 20S proteasome, we conducted cryo-electron tomography experiments and confirmed that particles stay close to the surface of amino-GO grids, rather than being located at the air-water interface. Moreover, addition of a polyethylene glycol (PEG) spacer between the amino group and the GO surface (amino-PEG-GO grid; **Figure 1B**), kept particles away from both the GO surface and the air-water interface, which may further reduce orientation bias and help preserve the sample integrity.

**Figure 1:**
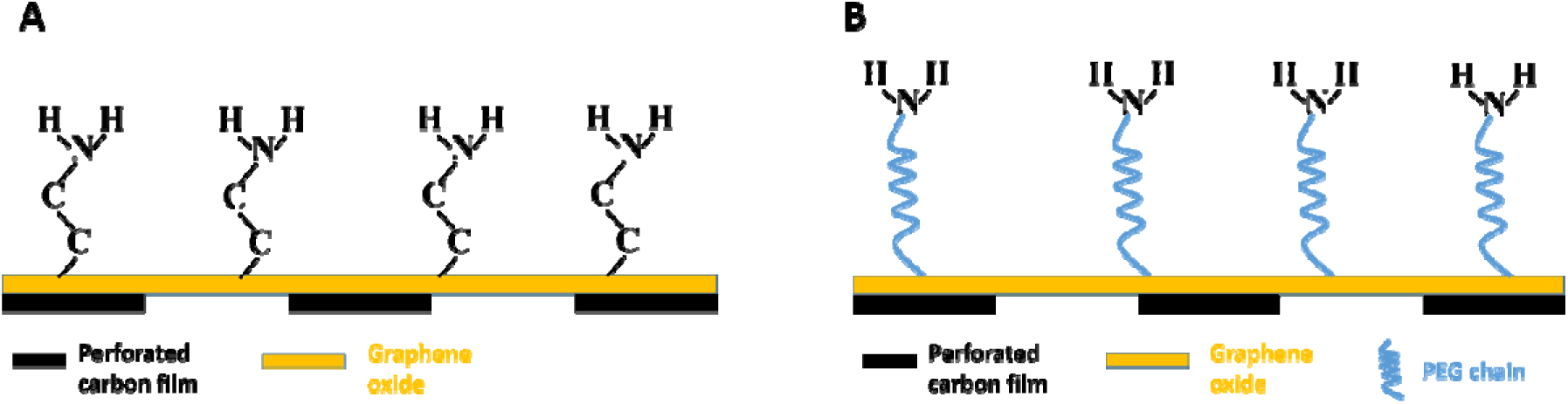
Schematic illustration of **(A)** amino-GO grid and **(B)** amino-PEG-GO grid assembly.

## 2. Materials and Methods

Synthesis of GO and deposition of GO onto Quantifoil EM grids were described in detail in our earlier work(Wang et al., 2019). Surface modification of GO was designed via the nucleophilic ring opening of epoxy groups by primary amines(Luo et al., 2016). GO grids were submerged in 30 ul of 10 mM ethylenediamine (Sigma-Aldrich E26266) solution in dimethyl sulfoxide (DMSO) and gently shaken for 5 hours. The grids were rinsed thoroughly with deionized (DI) water and ethanol sequentially and dried under ambient conditions. For Amino-PEG-GO grids, GO grids were submerged in 1 mM amine-PEG-amine (molecular weight 5000Da, Nanocs PG2-AM-5k) solution in DMSO and gently shaken overnight. After washing with DI water and ethanol, the amino-PEG-GO grids were air dried. Both kinds of grids should be stored dry at −20 °C and are sufficiently effective for months. Details of biological sample preparation and cryo-EM data collection can be found in the supplementary information.

## 3. Results and Discussions

### 3.1 GO coating on EM grid

As demonstrated in our previous work(Wang et al., 2019), GO deposition using the Langmuir-Blodgett method produced robust and satisfactory coating. It was our estimation that GO covers over 90% of the grid surface, with at least 40% being monolayer, about 40% bilayer and rest having three layers. GO regions having more than three layers are very rare. Two representative transmission electron microscope (TEM) images of GO coated gold Quantifoil EM grid at low magnifications are shown in **Figure 2**.

**Figure 2:**
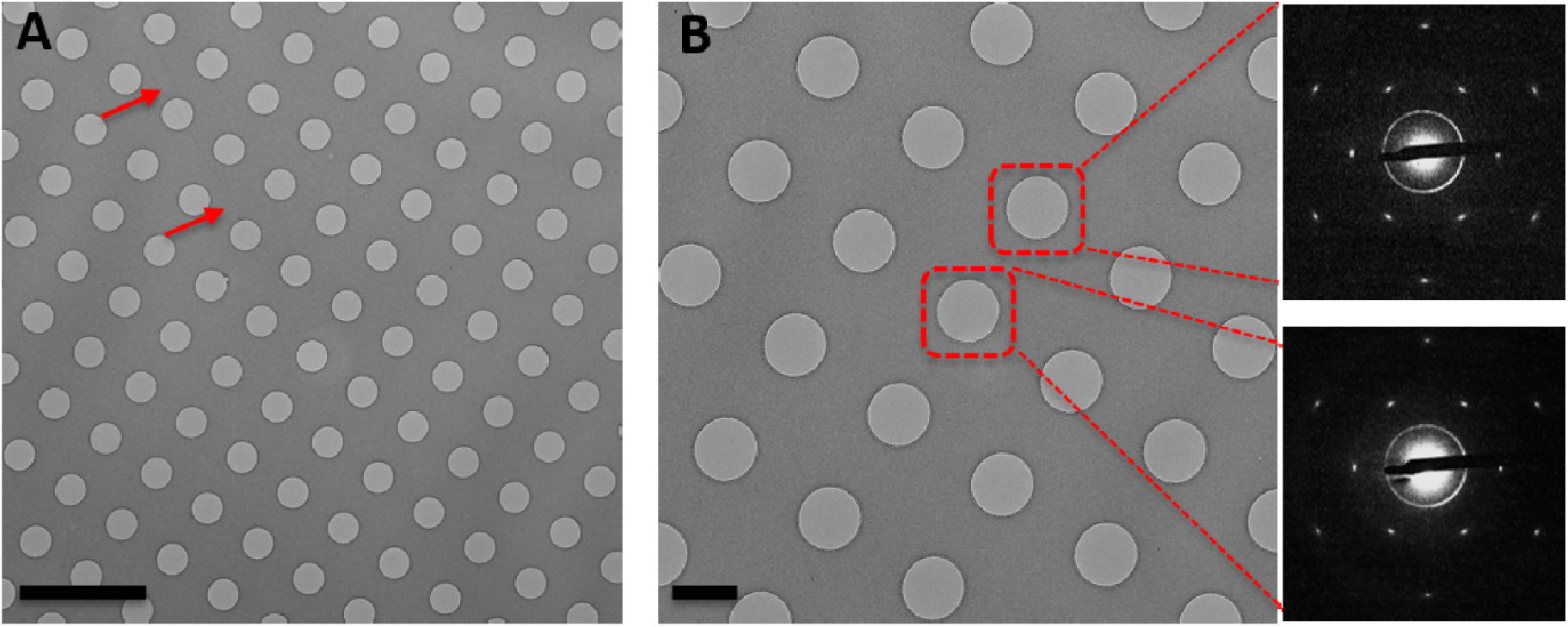
Transmission electron microscope (TEM) images of an EM grid covered by GO. **(A)** TEM image at 2500x displaying full coverage of GO over the square. The existence of GO is confirmed by wrinkles as indicated by the red arrows. Scale bar: 5 µm. **(B)** TEM image at 5000x showing GO coating of single layer. Selected area electron diffraction (SAED) images were taken from the holes marked by dashed square.

### 3.2 Improvement of protein distribution over the grid holes

We used the complex between bacterial Hsp90 and its client protein, bacterial ribosomal protein L2 (∼170kDa) to illustrate the performance of the amino-GO grid in protein distribution improvement. With Quantifoil holey carbon grids, no particles were observed inside the grid holes despite using complex concentrations as high as 10 µM (Figure 3A). The same sample showed ideal particle density when applied to amino-GO grids in concentrations as low as 250 nM, with excellent distribution, clearly recognizable particles and producing good 2D classes (Figure 3B and C).

**Figure 3.**
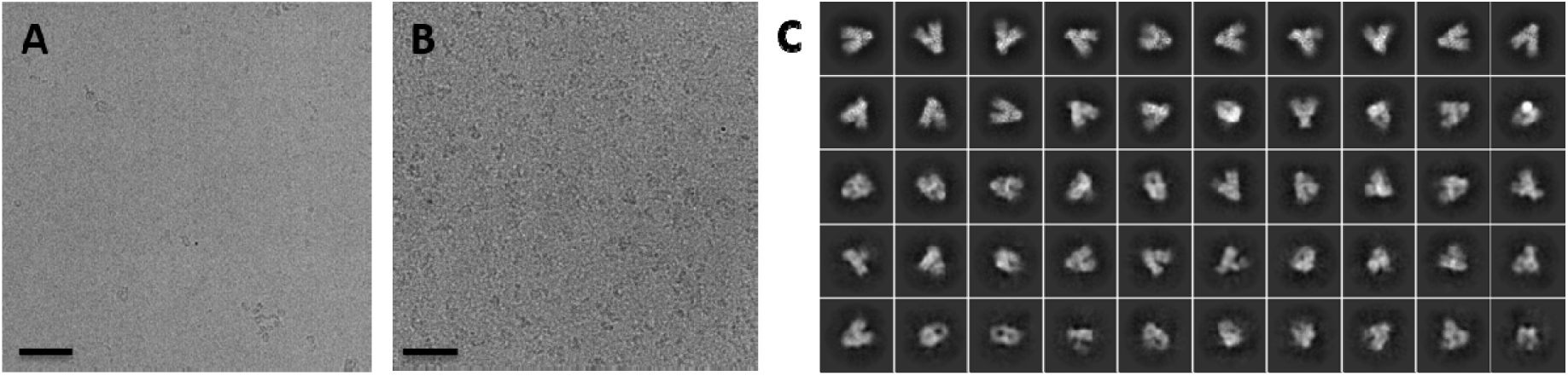
Cropped cryo-EM micrographs of the complex between bacterial Hsp90 and bacterial ribosomal protein L2 on a holey carbon grid **(A)** and on an amino-GO grid **(B)**. Scale bars: 50 nm. **(C)** Selected 2D class averages from the complex described in **(B)**. No particles were observable on a holey carbon grid with a protein concentration of 10 µM. By contrast, particle density was nearly ideal using an amino-GO grid with a protein concentration of 250 nM.

Another demonstration of improved particle distribution is evident when using a DNA origami sample. For this application, it was desired to have the DNA origami structure always land on its side, which requires the use of a support film on grid surface. Although a thin layer of amorphous carbon or GO film may both serve the purpose, we found that the DNA origami structures deform and make large aggregates on Quantifoil grids with thin amorphous carbon (**Figure 4A** and **D**) or GO films (**Figure 4B** and **E**). On the other hand, mono dispersed DNA origami structures were quite visible on amino-GO grids (**Figure 4C** and **F**), and remained well folded even though their large area flat surfaces were in contact with the film (**Figure 4G** and **H**).

**Figure 4.**
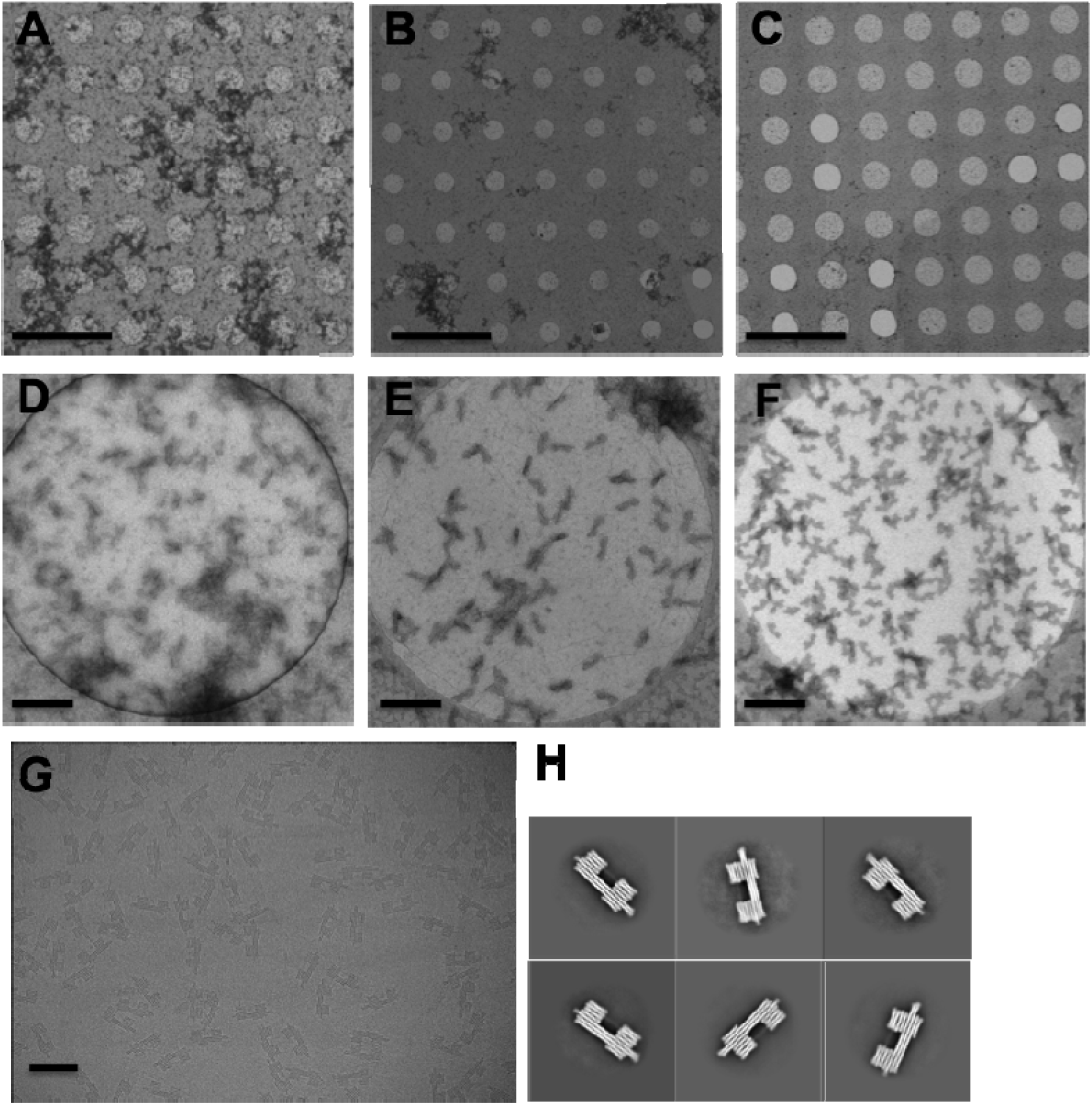
TEM images of DNA origami structures deposited on thin amorphous carbon coated grids (**A**) (**D**), GO grids (**B**) (**E**) and amino-GO grids (**C**) (**F**) from negative stain sampling. (**G**) A cryo-EM micrograph showing DNA origami structures observed on an amino-GO grid. (**H**) Selected 2D class averages of DNA origami structures from (**G**). Scale bars in **(A), (B)** and **(C)**: 5 μm. Scale bars in **(D), (E)** and **(F)**: 200 nm. Scale bar in (G): 100 nm. DNA origami structures aggregated severely on both thin amorphous carbon coated grids and unmodified GO coated grids, but dispersed well on amino-GO grids. For this sample, it was desired that all structures lay flat as observed with amino-GO grids (G).

### 3.3 Change of particle orientation

We tested the integrin αvβ8/L-TGF-β complex, which showed a set of strongly preferred orientations (predominantly side views) when frozen on traditional holey carbon grids at a concentration of 0.25 mg/ml. This resulted in 3D maps that were overestimated in resolution and highly “stretched” likely due to overfitting artifacts. The complex frozen on amino-GO grids (0.075 mg/ml) gave a wider distribution of orientations, providing more top, bottom, and ‘en face’ views than the holey carbon grids, as reflected on the orientation distribution map (**Figure 5**). Thus, using data from amino-GO grids was essential for obtaining a high quality map.

**Figure 5.**
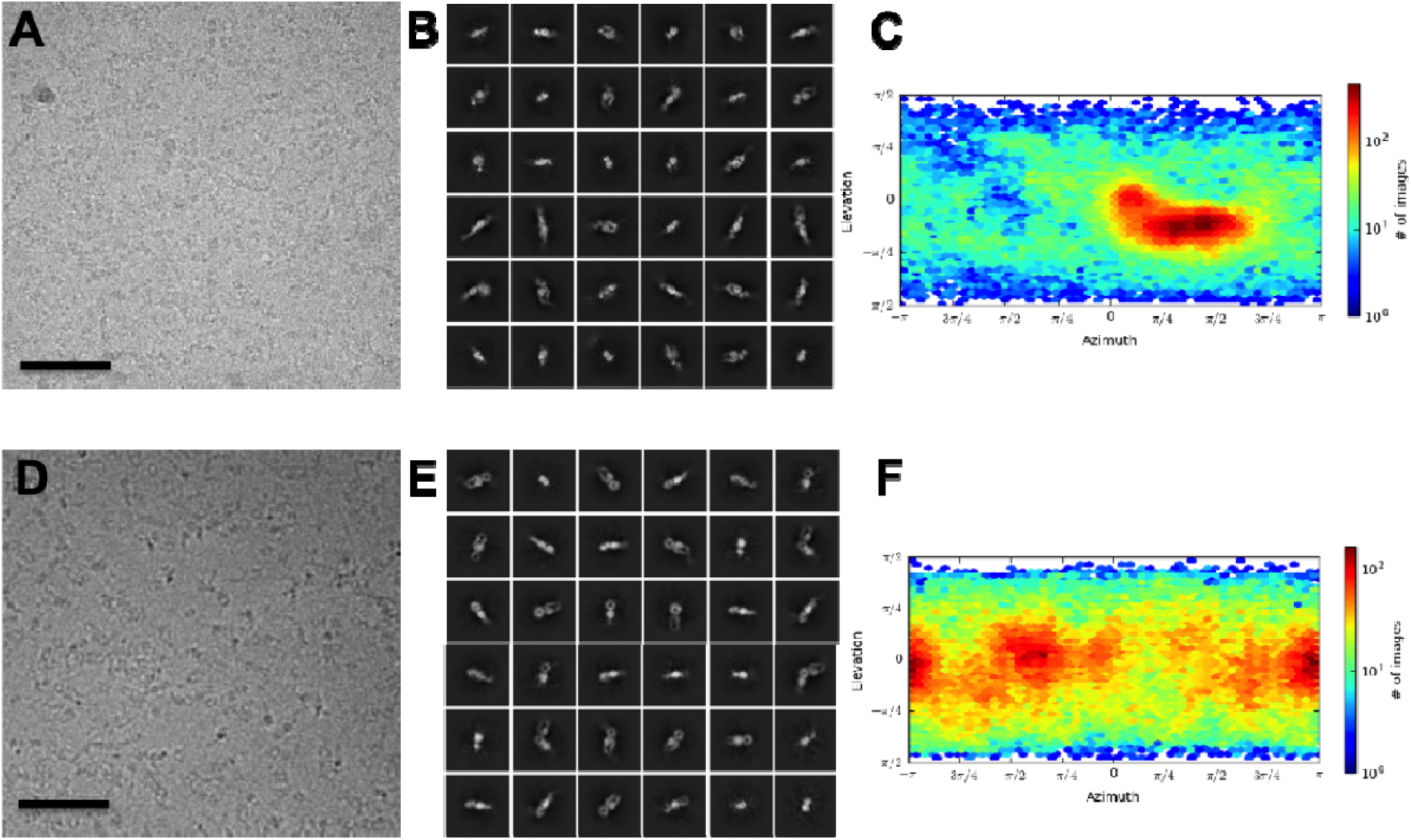
(**A**) Cropped cryo-EM micrograph, (**B**) selected 2D class averages and (**C**) orientation distribution of the αvβ8/L-TGF-β complex on traditional holey carbon grids. (**D**) Cryo-EM micrograph, (**E**) selected 2D class averages and (**F**) orientation distribution of the same αvβ8/L-TGF-β complex on an amino-GO grid. Scale bars: 50 nm. Particles exhibited predominately side views on holey carbon grids. On amino-GO grids, the orientation distribution improved dramatically.

Another project plagued by strongly preferred orientation was TRPA1, which adopted only top views on regular holey carbon grids, preventing 3D structure determination. In order to acquire side views necessary for calculating the 3D structure, the TRPA1 sample was prepared on amino-GO grids. As shown in Figure 6, TRPA1 particles adhered well to the grids, but this time exhibited a predominant side view orientation. A high-resolution 3D map of TRPA1 was then determined to ∼3.5 Å resolution by combining both orientations.

**Figure 6.**
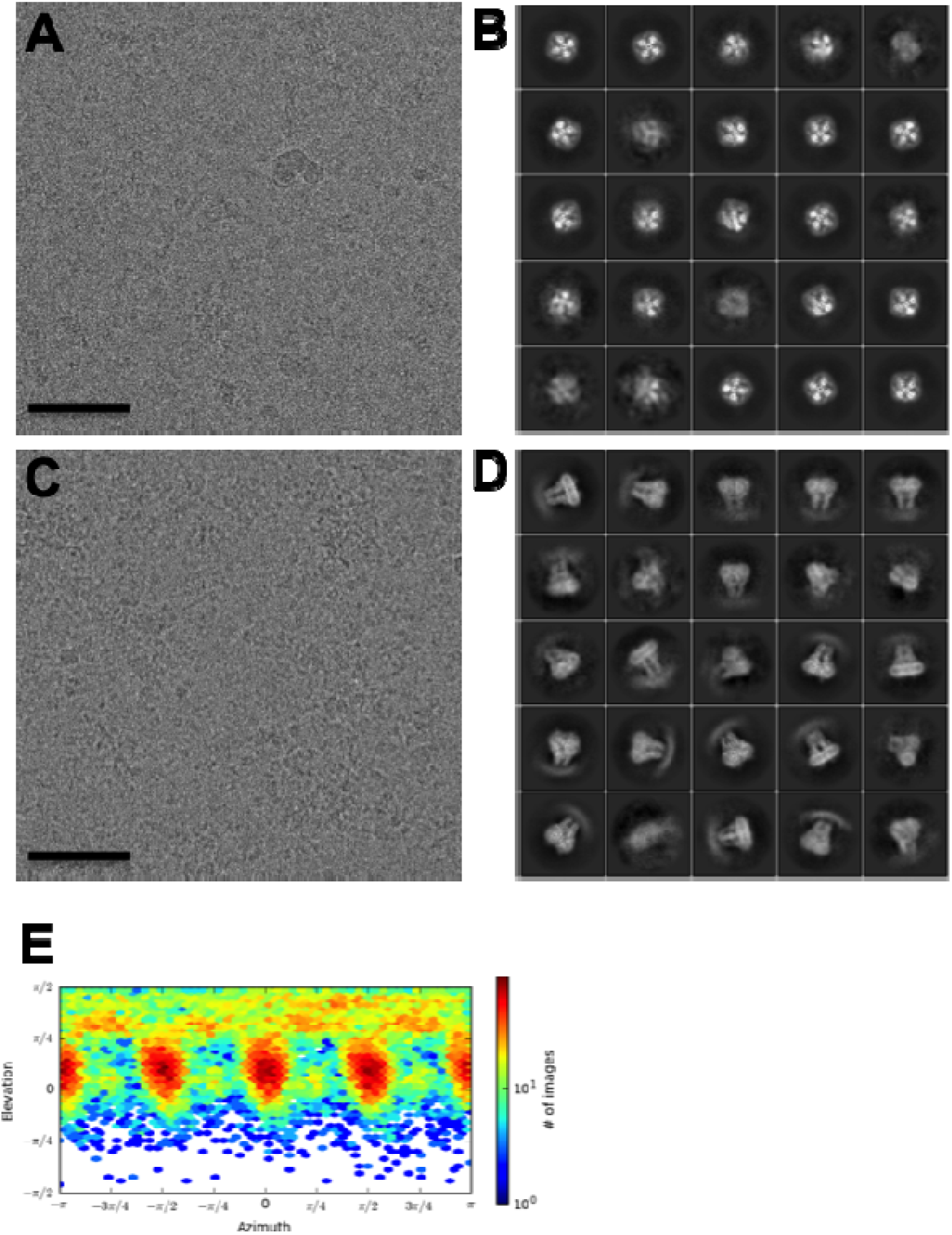
(**A**) Cropped cryo-EM micrograph and (**B**) selected 2D class averages of TRPA1 protein on a holey carbon grid. (**C**) Cropped cryo-EM micrograph and (**D**) selected 2D class averages of TRPA1 protein on an amino-GO grid. (**E**) Orientation distribution map of TRPA1 protein after combining data sets from both the regular holey carbon grid and the GO-amino grid. Scale bars: 50 nm. TRPA1 adopted only top views on holey carbon grids and only side views on amino-GO grids. Although neither is desirable by itself, the combination was quite successful.

**Figure 7:**
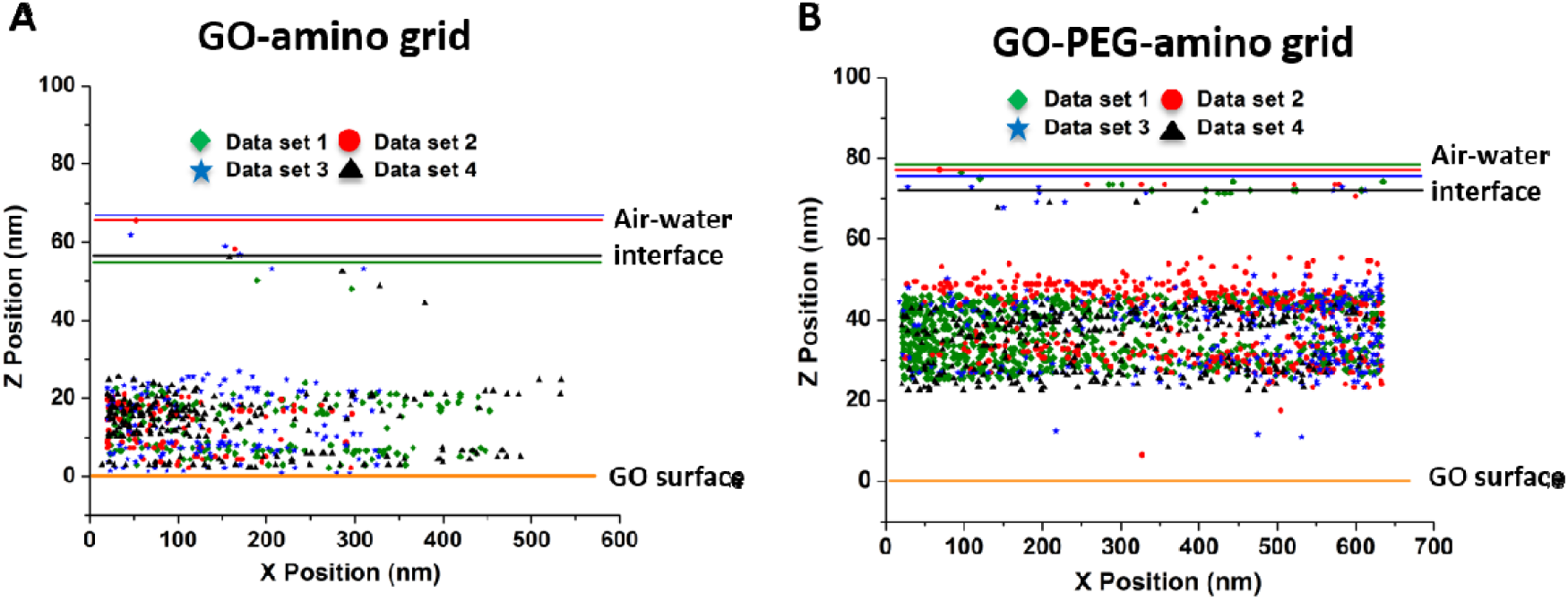
Determination of particle position from within the vitreous ice layer by tomographic analysis. Localization of 20S proteasome particles on (**A**) amino-GO grids and (**B**) amino-PEG-GO grids. Particle coordinates were collected from 4 data sets and represented by markers in different colors. The GO surface is set to zero for all data sets. The location of the air-water interfaces are illustrated with lines of color corresponding to each data set.

Considering that most particles dwell at the air-water interface with conventional holey carbon grids(Noble et al., 2018), it is reasonable for us to assume the orientation change was caused by keeping particles away from air-water interface. This was confirmed by tomographic analysis. Using the 20S proteasome, on the amino-GO grid the vast majority of protein particles were pulled away from the air-water interface, likely due to a weak electrostatic interaction between the primary amines on the GO and the protein. The particles stacked continuously spanning a distance of 20 nm with the bottom layer of particles around 5 nm from the GO surface. As 20S particle are cylinder shaped, the two bands of distances likely represent perpendicular orientations with some or most particles in near contact with the amino-GO surface. In order to keep particles away from both the air-water interface and the GO surface, we introduced a PEG spacer between the capping amino groups and the GO surface. As a result, with the same amount of protein applied, few particles were found to stick to the GO surface while particle stacking remained around 20-40 nm, again likely reflecting perpendicular orientations.

## 4. Conclusion

In summary, functionalizing GO grids with amino groups results in a support film for cryo-EM broadly capable of providing sample enrichment. The grid surface wettability as well as the protein distribution were greatly improved compared to bare GO grids. Moreover, the particle orientation could be changed due to the capability of the amino-GO grid of pulling particles away from the air-water interface. Notably, addition of a PEG spacer kept particles away from both the air-water interface and the GO surface, which would protect delicate samples from potential partial denaturation and aggregation, as well as minimizing orientation bias. It is also noteworthy that while we focused on amino modifications, GO grid surface functionalization conveniently extends to the coupling of a diverse array of moieties, i.e. carboxyl groups, thiol groups, secondary amines, DNA/RNA, hydrophilic polymers etc. to offer general affinity and adapt to a much broader range of proteins.

## Acknowledgements

We wish to thank Yanxin Liu and Eugene Palovcak for helpful discussions, Michael Braunfeld, Alex Myasnikov, David Bulkley for help and running the UCSF Advanced Cryo-Electron Microscopy Facility, and Matt Harrington for HPC support. TA holds Ruth L. Kirschstein NRSA Postdoctoral Fellowship F32GM119322. SMD is supported by NIH grant R35GM125027. Funding to DAA comes from a UCSF PBBR technology development grant, and NIH grants R35GM118099, U54CA209891, U01MH115747 and U19AI135990. YC is supported by NIH grants R01GM098672, 1R01HL134183, 1S10OD021741 and 1S10OD020054. DAA and YC are supported by the Howard Hughes Medical Institute, and the facility has received NIH instrumentation grants S10OD020054 and S10OD021741.

## Corresponding Author

Correspondence to David A. Agard

