## Supplementary material for "Amino and PEG-Amino Graphene Oxide Grids Enrich and Protect Samples for High-resolution Single Particle Cryo-electron Microscopy": Supplemntary information

^1^Department of Biochemistry & Biophysics and the Howard Hughes Medical Institute. University of California, San Francisco. San Francisco, CA 94143

^2^Department of Cellular and Molecular Pharmacology. University of California, San Francisco. San Francisco, CA 94158.

^ǂ^Feng Wang and Zanlin Yu contributed equally

**Complex Hsp90-L2**

The complex between full length bacterial Hsp90 (HtpG) and the bacterial ribosomal protein L2 was formed by incubating the two components at 10 µM Hsp90 dimer and 10 µM L2 for 20 minutes, crosslinking with 40 µM BS3 for 20 minutes and quenching the crosslinking reaction with 100 mM Tris-HCL. Proteins were expressed and purified as previously described(Genest et al., 2013).

Images were acquired on a Titan Krios microscope (Thermo Fisher Scientific) operating at 300 kV with a nominal magnification of 105,000x. Images were recorded using a Gatan K3 Summit detector (Gatan Company) with super resolution mode (0.424 Å/pix).

**DNA Origami**

The DNA origami structure was designed using Cadnano2 (Douglas et al., 2009). DNA staples were ordered from IDT. A custom M13 scaffold was prepared as described elsewhere(Douglas et al., 2009). DNA scaffold and the staples were stored in TE Buffer pH 8.0. The DNA origami structure was folded in a one pot reaction by mixing 10 nM DNA scaffold with 100 nM DNA staples mix in 20 mM MgCl_2_, 5 mM Tris-Base, 1 mM EDTA at pH8.0. After bringing the mix to 65 ˚C, it was then cooled down from 60 to 40 ˚C in 1˚C decrements, waiting at every temperature step 1 hour. Finally, temperature was brought to 20 ˚C and the sample was stored at 4 ˚C for later use. DNA origami structures were purified from excess staples using high concentration PEG-8000. First, unpurified DNA origami sample was brought to 7.5% w/v PEG-8000, 0.5M NaCl, 20 mM MgCl_2_, 5 mM Tris-Base, 1 mM EDTA at pH8.0 and it was spun at 25,000x g for 30 min at room temperature. The supernatant was aspirated gently and the opaque pellet was resuspended in DNA origami storage buffer (20 mM MgCl_2_, 5 mM Tris-Base, 1 mM EDTA at pH 8.0). PEG precipitation and resuspension was repeated one extra time to remove traces of excess staples and the purified DNA origami was stored in storage buffer at 4 ˚C.

Micrographs were collected on a Talos Arctica microscope (Thermo Fisher Scientific) operating at 200 kV with a K3 camera (Gatan), at a nominal magnification of 22,000x corresponding to a physical pixel size of 1.82 Å.

**Complex αvβ8/L**-**TGF-β**

Integrin and **L**-**TGF-β** were purified and a complex was formed as previously described (Campbell et al., *manuscript submitted for publication*). Data on the holey carbon grids were acquired on a Titan Krios microscope (Thermo Fisher Scientific) equipped with a K2 camera operated at 300 kV. Movies were recorded in super-resolution mode with a super-resolution pixel size of 0.67 Å and a nominal magnification of 105,000x. Data on the amino-GO grids were acquired on a Talos Arctica transmission electron microscope (Thermo Fisher Scientific) at 200 kV equipped with a Gatan K3 Summit direct detector. Movies were recorded in super-resolution mode with a super- resolution pixel size of 0.57 Å and a nominal magnification of 36,000x.

**TRPA1**

TRPA1 was purified as previously described(Paulsen et al., 2015). Data were collected on a Talos Arctica microscope (Thermo Fisher Scientific) equipped with a K3 camera operated at 200 kV, at a nominal magnification of 36,000x corresponding to a with a super-resolution pixel size of 0.57 Å.

**Tomogrphic analysis**

The electron cryo-tomography tilt series of 20S proteasome (0.1 mg/ml) were collected on a Talos Arctica microscope (Thermo Fisher Scientific) operating at 200 kV with a K3 camera. We used SerialEM(Mastronarde, 2005) to record movies in the counting mode with a pixel size of 1.82 Å. Data were acquired in a tilt range ±60^o^ with an interval of 3^o^, in two branches starting at 0^o^. The tilt series' were aligned using IMOD(Kremer et al., 1996) based on the gold fiducials embedded with the protein sample. Tomograms were reconstructed by TOMO3D(Agulleiro and Fernandez, 2015).
